# A universal and effective variational method for destriping: application to light-sheet microscopy, FIB-SEM and remote sensing images

**DOI:** 10.1101/2024.02.02.578531

**Authors:** Niklas Rottmayer, Claudia Redenbach, Florian O. Fahrbach

## Abstract

Stripe artifacts are a common problem for various imaging techniques such as light-sheet fluorescence microscopy (LSFM), electron microscopy and remote sensing. These artifacts are characterized by their elongated shapes and compromise image quality and impede further analysis. To address the primary challenge of removing the stripe artifacts while preserving the object structures we present here an improved variational method for stripe removal with intuitive parametrization. Comparison against previously published methods on images from LSFM, FIB-SEM and remote sensing by visual inspection and quantitative metrics demonstrates the superior capability of the approach. Based on synthetic LSFM data obtained by simulation of physical light-propagation we enriched our analysis by the comparison of processed images to ground truth data and quantitatively confirmed that our method outperforms existing solutions in terms of improved removal of artifacts and retention of image structures. The open availability of our solution <u>[1]</u>, the flexibility in handling variations in stripe orientation and thickness ensures its broad applicability across diverse imaging scenarios.

## 1. Introduction

Stripe artifacts are a common problem in several imaging techniques including light-sheet fluorescence microscopy (LSFM), focused ion beam scanning electron microscopy (FIB-SEM) and remote sensing. In general, these artifacts are characterised by highly elongated shapes of low width which point in a common direction. Their removal is required not only to improve visual quality but also to enable further analysis and image processing on the data. The examples shown in Fig.1 highlight the diversity of image structures and stripes encountered in practice. While stripes are periodic, thin, long and of low visibility in (c), much more severe artifacts of varying width, smaller length and higher intensity can be seen in (a).

**Fig. 1.**
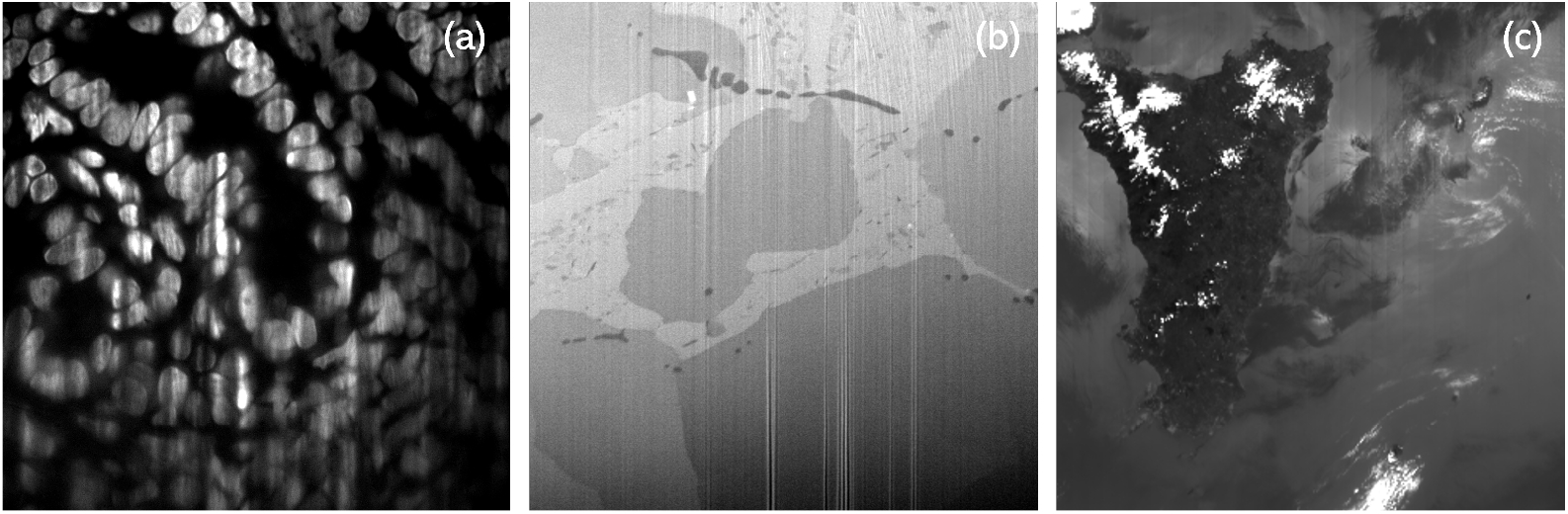
Stripe corruptions in cultivated mouse intestine cells imaged by LSFM (a), tin bronze cast alloy imaged by FIB-SEM (b) and Terra MODIS data from remote sensing (c).

There exists a large body of literature dealing with stripe removal including the recently published article by Ricci and colleagues [2] which provides a comprehensive overview of previous research on stripe removal in LSFM. Possible approaches are prevention [3–7], post-processing [8–12] and hybrid solutions [13, 14]. Preventing the formation of artifacts would be preferable but typically requires specialized hardware equipment or can limit the performance [15–17]. In contrast, post-processing is cheap and requires only computational power. Furthermore, corrupted images may already exist and specimens cannot be re-imaged, e.g., due to degradation and aging. For these reasons it is desirable to have powerful post-processing tools for stripe removal available.

In this article, we propose and explain a general solution for removing stripe artifacts. The results in this article quantitatively compare our method to previous approaches across images from LSFM, FIB-SEM and remote sensing. Our solution generalizes exceptionally well on all data tested. It is highly flexible thanks to intuitively adjustable parameters to accommodate for different appearances of structures and stripes. The source code is shared on GitHub [1] together with comprehensive guidance for application without further knowledge. The method which belongs to the class of variational methods is compared against selected state-of-the-art solutions from the same category and Fourier filtering methods. The performance is quantitatively evaluated with the help of synthetic LSFM data which is obtained by physically correct simulation of light transport with the Python package biobeam [18]. We include simulation data since it provides ground truth images which are unavailable for experimentally acquired images. This enables an objective assessment of performance using common quality metrics such as peak signal-to-noise ratio or the multi-scale structural similarity index measure (MS-SSIM). The evaluation is supplemented by a measure of stripe corruptions called “curtaining” proposed by Roldán [19] and the difference in line profiles.

In summary, we propose an improved solution for the removal of stripe artifacts, present results for a range of different imaging methods and, using several metrics, provide a thorough comparison with previously published solutions. Our work goes beyond previously published results as we demonstrate a rigorous analysis of results for different imaging methods and simulated data. Furthermore, we provide free access to well-documented code.

## 2. Methods

Most of the methods proposed for stripe removal belong to one of the two categories *Fourier filtering* and *variational methods*. We concentrate on these categories, as they generalize well to several imaging methods, variations in image structures and stripe appearance. Other approaches include average filtering [20, 21], histogram matching [22, 23], spline interpolation [24] and, recently, neural networks [25–28]. However, these methods are usually tailored for a specific appearance of images and stripes such that they are harder to transfer to other scenarios.

### 2.1. Variational Methods

Variational methods perform tasks such as denoising, segmentation, active contours [29] or destriping [30] by minimizing a task-specific convex objective function that is constructed by penalizing unwanted features of the image and stripes. Therefore, a minimizer of the function has desirable properties such as being free of stripe artifacts. Under mild assertions on the function, a unique minimizer exists and can be approximated using well-studied optimization algorithms, see [31].

Several variational methods for removing stripes were proposed in the past. The contributions [10, 32] represent stripes using elementary stripe patterns. Other propositions assume sparsity and a low-rank assumption on the stripe image [33–36]. Alternatively, [37] includes additional information from the Fourier domain. In the following, we introduce, motivate and explain our objective function which builds on previous work by Fitschen et al. [30]. A similar objective function to ours was also explored by Liu et al. [38] for remote sensing images. In contrast to previous implementations our objective function aims to effectively address stripes and retain image structures while providing a reasonable intensity profile.

#### 2.1.1. Optimization Problem

We assume that we observe a corrupted image *u*_0_ ∈ [0, 1]^*N*^ which can be decomposed as *u*_0_ = *u* + *s*, where *u* ∈ [0, 1]^*N*^ is the clean image and *s* ∈ [−1, 1]^*N*^ are the stripes. The dimensions of the image are *N* = *n*_*x*_ × *n*_*y*_ × *n*_*Z*_. Let ∇_*x*_, ∇_*y*_ and ∇_*Z*_ denote the directional difference operators for the three coordinate directions and assume stripes in *y*-direction. The objective function of variational methods is composed of a data term *D* (*u*) and noise term *N* (*s*), which independently penalize properties of the clean image and stripes. We propose

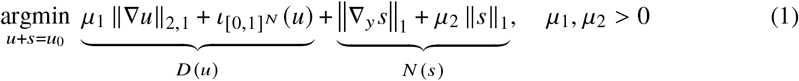

with indicator function 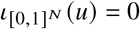 if *u* ∈ [0, 1]^*N*^ and ∞ otherwise and

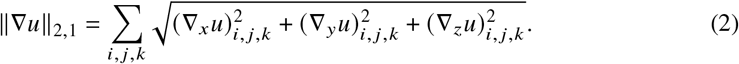

Term (2) is known as total variation [39] and encodes that clean images contain few strong edges. The choice of ||∇_*y*_ *s*||_1_ is based on the assumption that stripes have large differences orthogonal and small differences parallel to the stripe direction. Additionally, ||*s*||_1_ promotes sparsity in the stripe image and reflects that typically only a small part of the image is directly affected by artifacts. The indicator function 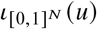 ensures that the output is in the same value range as the input. It prevents the need for brightness adjustments after processing and ensures non-negativity similar to natural intensities. The parameters *μ*_1_ and *μ*_2_ control the stripe removal, regularization and smoothness. They enable us to adapt our method to different conditions ranging from severe corruption to weak obscurities or ambiguities. For a discussion of the choice of *μ*_1_ and *μ*_2_ we refer to Supplement 1 Section 2. A basic intuition is given by:

- *μ*_1_ adjusts the strength of stripe removal. Larger values result in stronger removal but may affect stripe-like structures and yield smoothing.
- *μ*_2_ affects the precaution towards retaining image structures. Increasing *μ*_2_ enforces sparsity of the stripe image. Hence, larger values impair the removal, especially for non-stripe elements.
- The ratio of *μ*_1_ and *μ*_2_ is essential. Scaling both by an equal factor changes the amount of stripe removal while retaining the effect on image structures for the most part.

Optimization of (1) is done with the primal-dual gradient hybrid method with extrapolation of the dual variable (PDHGMp) [31]. The sequence generated by this iterative algorithm converges to the solution [31, 40]. To generalize (1) we propose to replace ∇_*Z*_ with 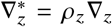 where *ρ*_*Z*_ ∈ = [0.1]. Setting *ρ*_*Z*_ = 0 reflects a 2D case while 0 < *ρ*_*Z*_ < 1 implies that information along the *Z*-direction is less reliable and coherent, e.g., due to lower resolution. For oblique stripes in direction *θ* we replace ∇_*y*_ *s* by ∇_*θ*_ *s* as suggested in [38]. Furthermore, simultaneous multi-directional destriping is possible by including a sum of penalization terms ||∇_*θi*_ *s* ||_1_ with different stripe directions *θ*_*i*_.

### 2.2 Fourier Filtering

The Fourier transform is a mathematical decomposition of an image into constituent frequencies. If the fundamental assumptions on stripes to be parallel and thin is fulfilled, stripe information is mostly encoded in frequency coefficients in a small band orthogonal to the stripe direction. In most cases, natural structures are dominated by low frequencies and concentrate near central coefficients. This decisive difference is exploited by Fourier filtering approaches by using masked filters [41–43], a decision-based algorithm [11] or a prior structural decomposition [8,9] to restrict tampering to stripe related coefficients. The latter approach reduces the influence on image structures the most. While wavelet-Fourier filtering [8] is commonly referenced, the approach by Liang et al. [9] using the non-subsampled contourlet transform (NSCT) [44] improves stripe and structure separation resulting in less interference during removal. We compare our method against a slightly enhanced version of this approach abbreviated by MDSR^+^. The method is explained in Supplement 1 Section 3.

## 3. Results

In this section, we evaluate and compare our method against previously published solutions from the category of Fourier filtering and variational methods. In particular, we consider the MDSR^+^, see Section 2.2, and the variational stationary noise remover (VSNR) [10]. For the MDSR^+^ we use the NSCT with 8 directions. The maximum possible depth, which depends on the image size, was set to either 4 or 5. The VSNR was used with three differently sized stripe patterns created from Gabor filters to be generally applicable. Our method (1) was used with *ρ*_*Z*_ = 0 as MDSR^+^ and VSNR were only available for 2D processing. The number of optimization steps taken for our method and VSNR were chosen to be 25000 to obtain a sufficient approximation of the optimum. The remaining parameters of the three stated methods were optimized via grid search, numerical and visual assessment. For the numerical assessment on synthetic data with ground truth we utilize common performance metrics: peak signal-to-noise ratio (PSNR) and the multi-scale structural similarity index measure (MS-SSIM) [45]. Furthermore, we consider the difference in line profiles of a selected area to the ground truth using the euclidean norm. Additionally, we use the curtaining metric proposed by Roldán [19]. In contrast to the other metrics, it exclusively measures stripe corruptions based on the ideas of Fourier filtering methods, see Section 2.2. We report the absolute difference between the curtaining metrics of the processed image and the ground truth. This choice adjusts for naturally occurring stripe-like patterns in the image structures and treats over- and under-performance alike.

### 3.1. Synthetic Data

Fig.2 shows results on a synthetic drosophila-like embryo model inspired by [18] and obtained by physically accurate modeling of light propagation in LSFM imaging, see Supplement 1 Section 1 and [46]. Major differences between the methods can be spotted. While MDSR^+^ reduces the stripes significantly, larger artifacts and thin oblique stripes remain visible in the outcome. The latter become only visible at closer inspection. In comparison, VSNR is able to reduce wider artifacts better with only weak remnants. However, fine oblique artifacts are entirely unaffected by the removal. Only our method removes all stripes including the trails with only faint artifacts remaining in particular at the right edge of the body. The line plot supports our findings. MDSR^+^ shrinks the profile but is unable to adjust wider artefacts correctly. On the other hand, the line profile of VSNR has less extremes but contains small perturbations reflecting the struggle with fine artifacts. Our method is the only one which has both a significant reduction in stripes and a flat profile. The numerical values presented in Table 1 further confirm prior observations with our method outperforming the others in all metrics.

**Fig. 2.**
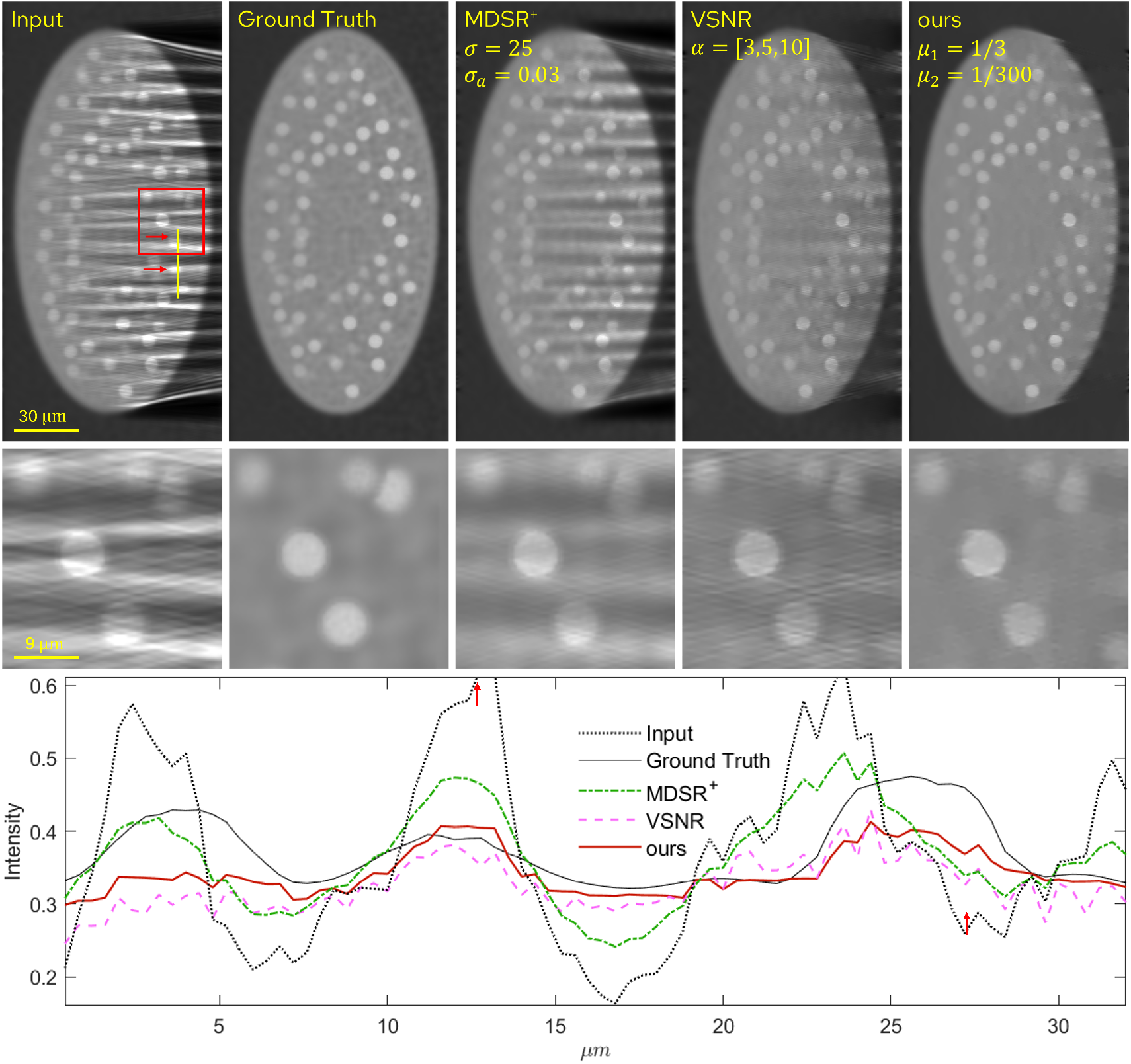
Destriping results and line profiles of simulated LSFM data of a synthetic drosophila-like embryo model.

**Table 1.** Quantitative evaluation of results on synthetic data from Fig.2. Arrows indicate better performance. Percentage improvements from the input are given in green.

| Metric | Input | Ground Truth | MDSR<sup>+</sup> | VSNR | ours |
| --- | --- | --- | --- | --- | --- |
| PSNR↑ | 24.25±0% | ∞ | 29.02<sup>+20%</sup> | 31.06<sup>+28%</sup> | 35.65<sup>+47%</sup> |
| MS-SSIM↑ | 0.901±0% | 1 | 0.940<sup>+4%</sup> | 0.919<sup>+2%</sup> | 0.969<sup>+7.5%</sup> |
| $\partial$Curtaining↓ | 0.184±0% | 0 | 0.056<sup>-70%</sup> | 0.074<sup>-60%</sup> | 0.036<sup>-80%</sup> |
| Line difference↓ | 1.055±0% | 0 | 0.523<sup>-50%</sup> | 0.589<sup>-44%</sup> | 0.389<sup>-63%</sup> |

### 3.2 Real Data

In Fig.3 we show results obtained by processing real data from LSFM, FIB-SEM and remote sensing. We observe similar behaviour to the synthetic data with VSNR leaving very thin remnants and MDSR^+^ having insufficient reduction. The LSFM image highlights the capabilities of variational methods in general. While MDSR^+^ only reduces stripes, both VSNR and our method show near perfect on par performance. However, the FIB-SEM and remote sensing images reveal short-comings of VSNR which leaves very thin stripes in the result. MDSR^+^ even introduces severe image artifacts in areas of high contrast in the remote sensing image despite performing well in low contrast regions. On the other hand, our method delivers visually perfect results for both FIB-SEM and remote sensing in the sense that stripes are entirely removed and image structures remain intact.

## 4. Discussion

The results shown in Section 3 demonstrate that our method produces consistently good results in terms of stripe removal quality and retention of image details while offering intuitive adjustability through its two weighting parameters *μ*_1_ and *μ*_2_. Results shown in Section 2 in the Supplement illustrate the impact of these parameters. Also, we provide information on their ideal choice. The consistency across all shown settings can be attributed to the non-restrictive formulation of its objective function with only rough assumptions on properties of the image and stripes. Therefore, deviations in stripe direction and thickness remain adequately penalized such that a general application for stripe removal is provided. However, limitations exist. In particular, the alignment of image structures with the stripe direction should be avoided since both will become affected by the removal, see Supplement 1 Section 4 for more details.

The insufficient removal by MDSR^+^ for more sophisticated stripes is due to a large overlap in area of stripe and image related frequency coefficients when structures and stripes live on similar scales or stripe directions deviate from the assumption. Depending on the choice of parameters, either insufficient stripe removal or severe modification of image structures are the result. We deem the prior to be preferential since the latter renders the outcome unusable for any analysis or visualisation which is why we only showed these results. The introduction of artifacts in the remote sensing image is surprising as the setting reflects an ideal scenario for this approach with thin, long and periodic stripes. However, the high contrast in some areas produces unforeseen problems by filling separating spaces with smooth stripe-like artifacts.

In contrast, VSNR performs overall well but has problems removing thin or oblique stripes. This behaviour arises since the considered stripe patterns do not sufficiently reflect these particular instances of stripes. We see potential of VSNR to improve using more image specific patterns as demonstrated by the results in the top row of Fig.3 which were achieved by using optimized patterns. However, results are with less optimized patterns are inferior and we argue that optimizing patterns and corresponding parameters requires considerable expertise, is time consuming and impractical for a general use.

**Fig. 3.**
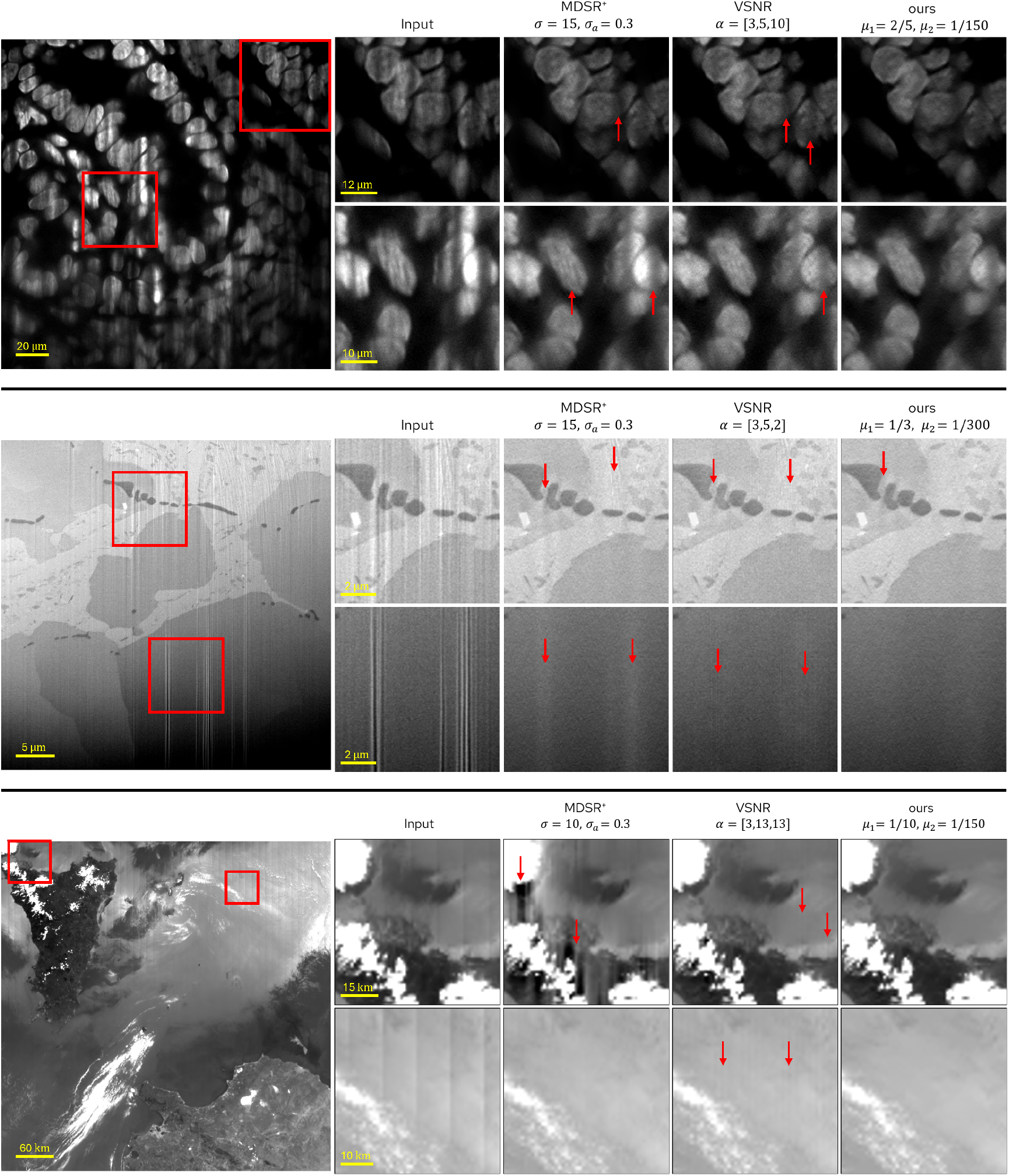
Stripe removal results on real data acquired by LSFM (top), FIB-SEM (middle) and remote sensing (bottom). The red arrows highlight areas of particular interest.

For our analysis we supplement real data with synthetic LSFM data. We used a highly accurate simulation method which yields realistic, irregular stripe artifacts. It is important to note that realistic models of the refractive index distribution within the sample are crucial. Therefore, considerable effort and rigour was applied to creating a realistic model following [18]. While simulations have been employed for the assessment of LSFM performance such as [46], so far stripe removal algorithms have not been evaluated based on the synthetic data despite the clear benefits: the objective assessment by comparison to the ground truth obtained by “turning off” the cause of the stripe artifacts yielding ground truth unperturbed images for comparison. This approach allowed the use of quality metrics such as PSNR and thus a more accurate assessment of quality compared to the previously employed approach of superimposing a clean image with independent stripes [9] or relying purely on visual assessment.

Roldáns curtaining metric [19] complements previously established metrics well for assessing quality on the synthetic data. Since it is specifically designed to measure the amount of stripe corruptions in an image it is more robust against other difference between the image and its ground truth, e.g., different levels of brightness. We did not apply the metric directly but used the absolute difference with the ground truth. We have found that a poor choice of parameters leading to an overly aggressive stripe removal often results in a seemingly better curtaining value, although the methods perform poorly in all other relevant aspects, such as structure preservation. By adjusting for naturally occurring imperfections in the structures using the ground truth, the metric captures improvements in image quality more accurately and is in line with visual inspection and the other metrics.

The results shown in this article were processed as 2D images. This approach does not reduce the significance of our findings but simplified visualization and comparison with imaging methods that deliver 2D images like FIB-SEM and remote sensing and processing algorithms that work only in 2D. However, we emphasize that our method is able to process 3D images as well. Importantly, image stacks are not processed slice by slice but fully in 3D. This improves consistency between slices and prevents possible errors in image structures that result from ignoring relations in the stack direction. However, in many cases the impact is limited due to inferior axial resolution which impairs the separation between image structures and stripes on this axis.

## 5. Conclusion

In this work, we have demonstrated the potential of an improved variational method using a specially adapted objective function as a general solution for removing stripe artifacts. Results for a variety of image structures and different imaging methods together with quantitative assessment demonstrated the potential of the method. An important aspect for the adoption of the method which we share openly [1] is the correspondence of the parameters to intuitive image properties enabling non-specialist users to readily achieve optimized destriping and preservation of image structures. Furthermore, we substantiate that it is highly effective in removing stripes and exceeds the performance of previously published solutions. The generality of the approach is backed by using images from three different imaging methods: LSFM, FIB-SEM and remote sensing. A quantitative comparison on synthetic LSFM data using several quality metrics helped to better understand the capabilities and limitations of the stripe removal methods. It confirmed that our solution provides best destriping while retaining image structures which coincides with visual assessment of real images.

## Supplemental document

See Supplement 1 for supporting content.

## Funding

Federal Ministry of Education and Research (BMBF), Project: Synthetic Data for Machine Learning Segmentation of Highly Porous Structures from FIB-SEM Nano-tomographic Data (poSt), Funding number: 01IS21054A

## Acknowledgments

We thank Martin Weigert for guidance on installing biobeam and its application for modelling LSFM imaging as well as Michael Engstler for providing the FIB-SEM image data. Furthermore, we want to thank Diego Roldán for providing and explaining his curtaining metric and Jannik Reiser for his help with data processing. Lastly, we thank Nazar Oleksiievets, Sebastian Bundschuh, Stephan Preibisch, Michael Innerberger and Panagiotis Symvoulidis for using our code and providing valuable feedback.

## Disclosures

The authors declare no conflicts of interest.

## Data Availability Statement

The methods presented are available in a GitHub repository [1] which includes code for all used methods and scripts for processing images. Furthermore, a Jupyter-notebook for generating synthetic LSFM data and a txt-file for creating the Python environment on Windows 11 are provided.

## Notes

https://github.com/NiklasRottmayer/General-Stripe-Removal.git

### Competing Interest Statement

The authors have declared no competing interest.

### Summary of Updates

Minor updates to the text and abstract

