## Supplement 1 for "A universal and effective variational method for destriping: application to light-sheet microscopy, FIB-SEM and remote sensing images"

### CONTENTS


1 LSFM Simulation 1

2 Parameter Selection 2

3 $\text{MDSR}^+$ 2

4 Additional Results 4


### 1. LSFM SIMULATION

Creating synthetic LSFM images requires the simulation of light propagation through media to capture effects such as scattering and absorption correctly. We achieve this using the python package *biobeam* [1] and by modelling geometric structures via a refractive index distribution (rid) and fluorescence distribution (fld). The rid is denoted by $\mathbf{n} \in \mathbb{C}^{n_x \times n_y \times n_z}$ containing complex valued refractive indices. The real part depicts the classical refractive index describing scattering behaviour. The imaginary part is the absorption coefficient and yields effects such as light attenuation. We refer to [2] for typical refractive indices for different parts of a biological sample. Ground truth images were generated by setting $n \equiv n_0$ to be constant. Typically, $n_0 = 1.33$ which is approximately the refractive index of water, the main constituent of biological cells. The fld $f \in \mathbb{R}_{\geq 0}^{n_x \times n_y \times n_z}$ contains non-negative factors which represent the proportion of re-emitted light by fluorescence when subjected to illumination. This indirectly models the real process of staining a sample with a fluorescent marker which are specialized proteins that emit light when illuminated at specific wavelengths. Besides geometrical distributions we select the wavelength of illumination light $\lambda$, the numerical apertures of illumination and detection optics $\text{NA}_{\text{ill}}$ and $\text{NA}_{\text{det}}$ and the voxel size. Furthermore, we set the illumination direction along the $y$-axis. Afterwards an image can be simulated by the following steps:

(i) **Illumination:** A complex valued $x$-$z$ cross-section of a cylindrical light-sheet is generated from $\text{NA}_{\text{ill}}$ and $\lambda$. With wave-optical propagation as described in [3], the cross-section is propagated through the sample. This yields the illuminated volume $I_{\text{ill}}$.

(ii) **Fluorescence:** The amount of light re-emitted through fluorescence is obtained via $I_{\text{fl}} = I_{\text{ill}} \odot f$ with element-wise multiplication $\odot$. It represents the 'fluorescence response' to the illumination.

(iii) **Acquisition:** The 2D image of the section in focus is acquired by convolution with the detection point-spread function (PSF) $h_{\text{det}}$ obtained from $\text{NA}_{\text{det}}$, i.e.

$$I_{\text{det}} = (I_{\text{fl}} * h_{\text{det}})_{z=\bar{z}}. \tag{S1}$$

(iv) **3D Imaging:** Repeating steps (i)-(iii) for different focal planes a stack of images is acquired and combined.

The essential parts of the simulation procedure are visualized in Fig. S1. For model generation in this paper, we sample non-overlapping spheres as foreground and combine the structures with 3D Perlin noise. Refractive indices were chosen based on reference values given in [2]. The values for the fld were chosen to produce visually appealing images that look similar to real LSFM images.

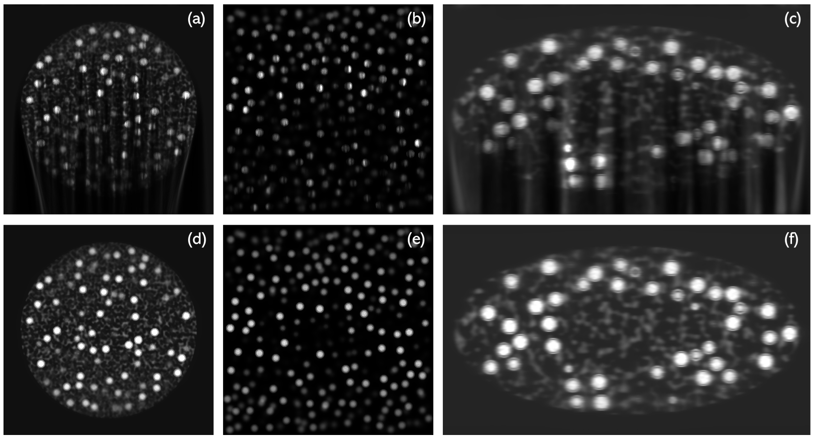


**Fig. S1.** Simulated images (a-c) and ground truths (d-f) for different simulation examples. Cell cluster (a,d), dispersed cells (b,e) and embryo (c,f).

### 2. PARAMETER SELECTION

For our proposed method, we want to give advice on how parameters should be selected and tuned. Fig. S2 displays a range of parameter pairs for $\mu_1$ and $\mu_2$ with the corresponding results. For this particular test image, the effects of $\mu_1$ and $\mu_2$ become most apparent in the extreme cases, e.g., the columns $\mu_2 = 1/600$ and $\mu_2 = 2/75$ showcase that $\mu_1$ controls the strength of reduction. The larger the value of $\mu_1$, the stronger the reduction of stripes but also the greater the risk of affecting and removing image structures. Conversely, the rows $\mu_1 = 1/10$ and $\mu_1 = 1/2$ highlight the effect of $\mu_2$. The larger $\mu_2$, the weaker the stripe reduction but it is less likely to affect image structures. From our experience of several examples, scaling both parameters by a comparable factor allows to address less ideal stripes, which e.g. deviate in direction or thickness. The ability to increase the parameters is limited by the impact on image structures. In combination, the parameters allow to adjust the strength of stripe removal, reduce smoothing artifacts and prevent the modification of image structures for different imaging methods and imaged subjects.

Depending on the severity of stripe corruptions and the appearance of stripes, different choices of $\mu_1$ and $\mu_2$ are advised and optimal. In general, $\mu_1 = 1/3$ and $\mu_2 = 1/300$ is a good starting point and the intervals $\mu_1 \in [0.1, 0.5]$ and $\mu_2 \in [0.0016, 0.017]$ were never exceeded. If stripe artifacts are less severe, $\mu_1 = 1/6$ or $\mu_1 = 7/30$ with $\mu_2 = 1/300$ is sufficient. Persistent stripes which are e.g. short and similar to structures benefit from using a stronger removal via $\mu_1 = 0.5, \mu_2 = 0.017$ or $\mu_1 = 0.4, \mu_2 = 1/150$.

### 3. MDSR$^+$

The multi-directional stripe remover (MDSR) [4] is a Fourier filtering-based stripe removal method. Its fundamental concept coincides with masked filter approaches [5–7] or the decision-based algorithm in [8]. It closely follows the ideas proposed by Münch et al. [9] of combining a structural decomposition with selected filtering of stripe-related subimages in the Fourier domain. The initial image decomposition of the MDSR is the non-subsampled contourlet transformation (NSCT) [10] which decomposes an image $u_0$ by scale and direction using a pyramidal filter bank. Afterwards, subimages $u_0^{(i)}$ attributed to directions close to the stripe direction are further processed with a typical Fourier filtering procedure:

$$\hat{u}_0^{(i)} = \mathcal{F}^{-1}\left[\left(1 - \exp\left(-\frac{y^2}{2\sigma_i^2}\right)\right) \odot \mathcal{F}(u_0^{(i)})\right], \quad \sigma_i = \sigma \cdot \exp -\frac{\theta_i^2}{2\sigma_a^2}$$

with element-wise multiplication $\odot$ where $y$ is the vertical distance to the image center, $\theta_i$ the angle of the subimage to the stripe direction, $\sigma$ the Gaussian decay parameter and $\sigma_a$ a decay factor for reduced impact for images further from the stripe direction. We refer to this method as MDSR$^+$ to differentiate it from the original proposition [4] which filtered all subimages. In our instance, we restrict filtering to directions $\theta \in [-\pi/4, \pi/4]$ from the stripe direction following the results shown in Fig. S3 which indicate that this significantly reduces artifacting.

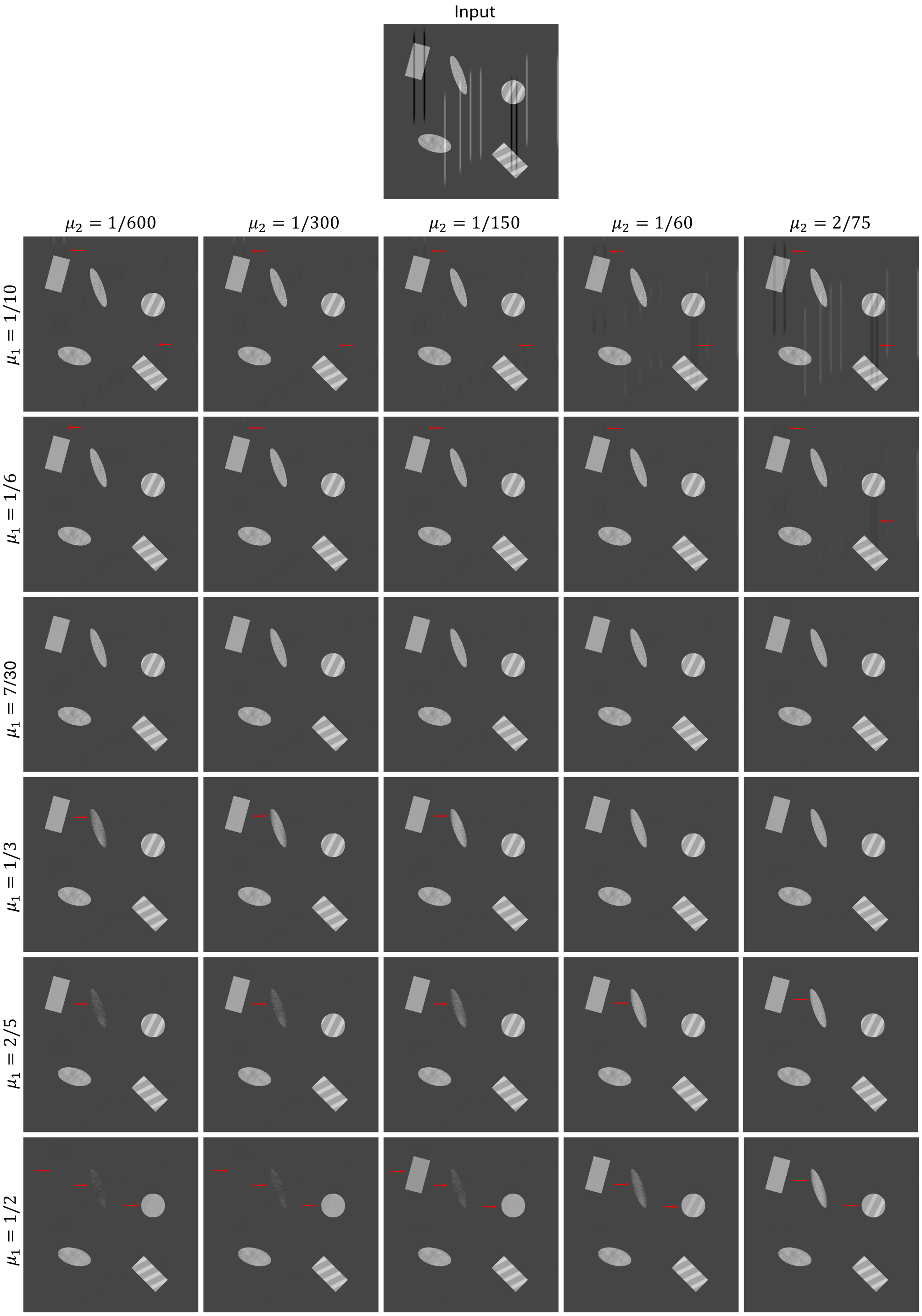


**Fig. S2.** Results of our method for different combination of parameters. The red arrows indicate image artifacts.

The MDSR$^+$ depends on $n_{\text{dir}}, n_{\text{dec}}, \sigma$ and $\sigma_a$. However, $n_{\text{dir}} = 8$ yields a sufficient directional decomposition and $n_{\text{dec}}$ must only be large enough such that stripes are captured by the NSCT. In our experiments, $n_{\text{dec}} \in \{4, 5, 6\}$ sufficed depending on the image size. The main parameter of this method is $\sigma$ controlling the strength of damping. An adjustment to $\sigma_a$ may also improve performance if stripe artifacts are not ideally vertical.

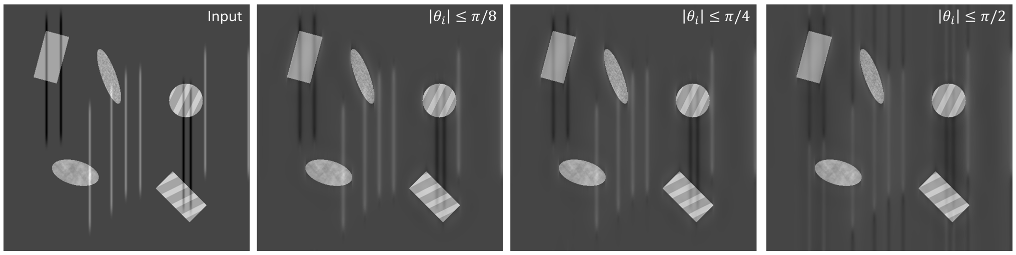


**Fig. S3.** MDSR$^+$ results of filtering in restricted directions using $\sigma = 10$ and $\sigma_a = 0.3$. The right image is the result of the original MDSR.

### 4. ADDITIONAL RESULTS

Fig. S4, S5 and S6 show several results of our method on data from LSFM and FIB-SEM. The results highlight the universal applicability of our method to almost all stripe corrupted images. In all shown instances the removal of stripes is performed with near perfection.

Fig. S4 highlights advantages of processing in 3D compared to 2D with increased consistency in the side and top view while showing equal performance in the front view. The stripes visible in the side view typically correspond to stripe artifacts in the front view. However, some are caused by variations in the brightness levels between the slices caused by the imaging procedure. The variations in brightness are also visible as horizontal stripes in the top view and remain entirely visible when processing 2D, i.e. slice-wise. In contrast, processing in 3D results in the removal of those artifacts since they are detected as planar stripes which appear only in individual slices and are not consistent with image structures.

The bottom row in Fig S6 reveals that when structures and stripes align, both will be reduced and removed by our method. This limitation is expected since the objective function of our variational method only distinguishes stripes and structures indirectly through their feature set. However, this limitation can sometimes be overcome by reducing the strength of removal, i.e., by reducing $\mu_1$ and changing $\mu_2$. This results in wider artifacts to remain strongly visible, but fine stripes to be removed while image structures are retained. This workaround can be feasible depending on the purpose of the destriped data. Nonetheless, we suggest to plan imaging of materials and structures accordingly such that the direction of stripes differs from the direction of structures. Another limitation shows up in the the top row of Fig S6 where severe corruptions are contained that suppress image information entirely. In these areas, only the level of brightness is corrected but previously lost information such as texture cannot be reconstructed. This problem can only be resolved by preventing severe artifacts from forming, see [11] for more details on prevention methods.

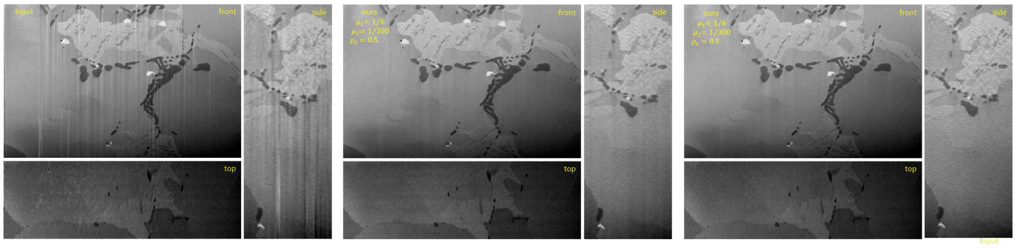


**Fig. S4.** Comparison of 2D and 3D destriping using our method on FIB-SEM images of a tin bronze cast alloy.

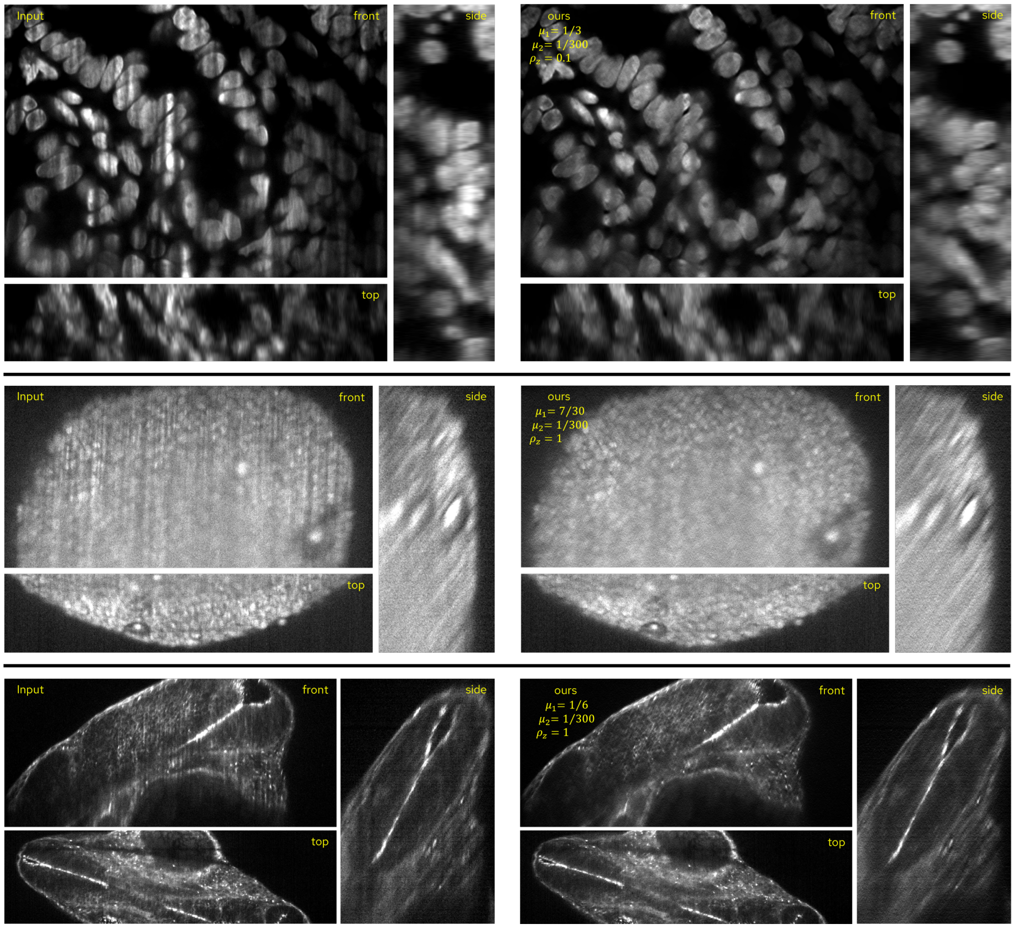


**Fig. S5.** Stripe removal results of our method on LSFM images of cultivated mouse intestine cells (top), a cluster of HeLa cells (middle) and the embryo of a zebra-fish larva (bottom).

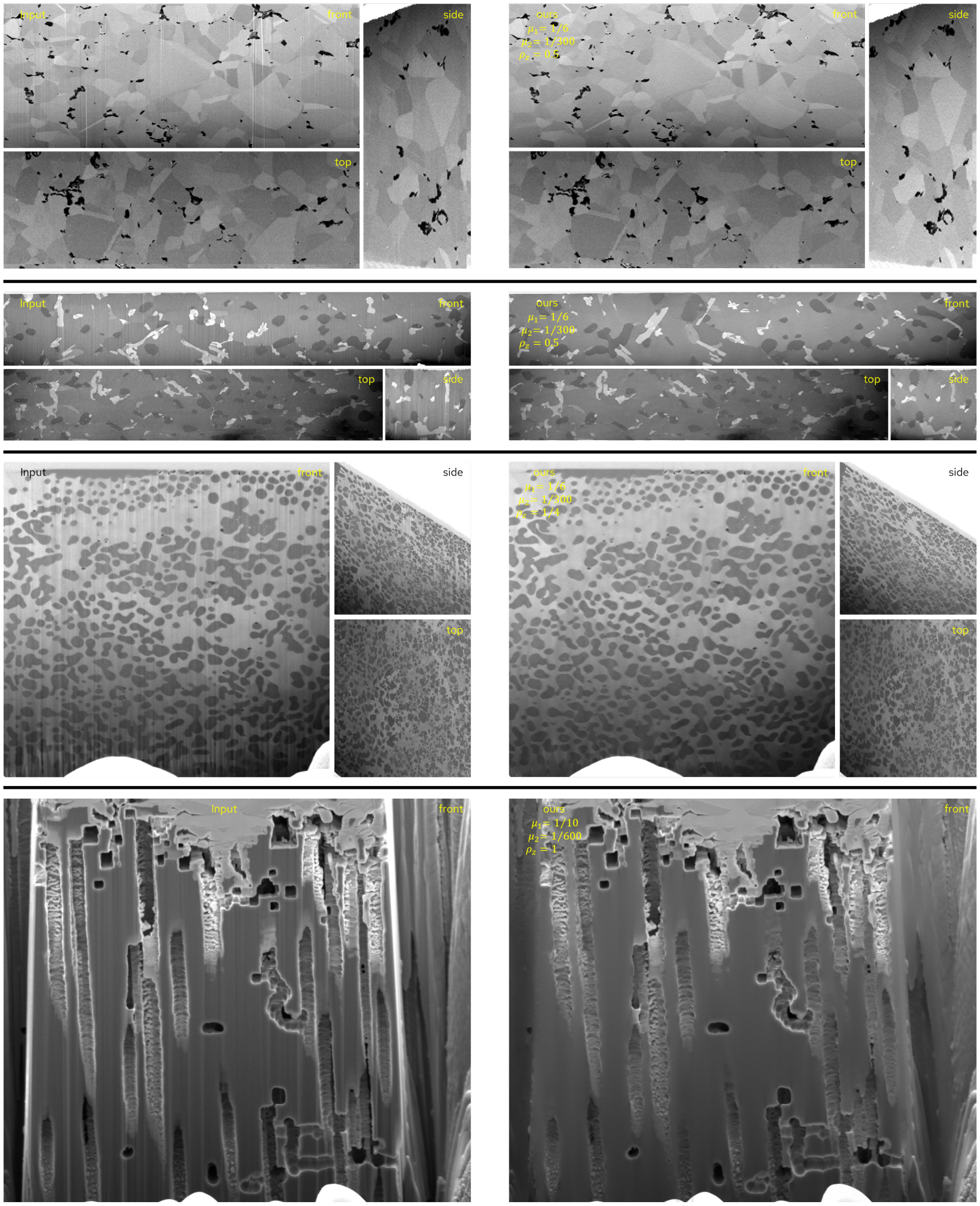


**Fig. S6.** Stripe removal results of our method on FIB-SEM images of carbon nanotube reinforced metal matrix composite (top), AlSi13 (2. row), CuCr (3. row) and porous aluminum (bottom).
